# Sequencing complete plasmids on Oxford Nanopore Technology Sequencers using *R2C2* and *Chopper*

**DOI:** 10.1101/2025.01.16.633418

**Authors:** Kayla D. Schimke, Christopher Vollmers

## Abstract

Plasmids are ubiquitous tools in molecular biology which are used for a large variety of experiments within academic and commercial labs.

Both new and old plasmids have to undergo sequencing-based analysis to determine whether or not they are functional, i.e. contain the correct insert in the correct backbone. While traditional Sanger sequencing based analysis was most often limited to the inserts, new high-throughput sequencing based methods and services can now provide the complete sequence of a plasmid. Currently available methods and services vary in throughput and cost.

Here, we adapted the Oxford Nanopore Technologies-based R2C2 sequencing method to - rapidly and at low cost - sequence complete plasmids, either individually or in a pool. We also developed an analysis pipeline, Chopper, that produces full-length plasmid sequences. We tested our workflow with commonly used plasmids we ordered from Addgene and produced highly accurate sequences for each plasmid from both their individual and pooled sequencing runs.

## INTRODUCTION

Plasmids are fundamental tools in molecular biology serving as vectors for recombinant DNA. Following their assembly, as well as periodically after, their DNA sequence has to be validated to ensure they are functional and retain their functionality over time.

Traditionally, plasmid sequencing has been performed using Sanger sequencing and has focused on the specific inserts in the plasmids, like the coding sequence of a protein, leaving the often reused backbones of the plasmids - promoters, origins of replication, antibiotic resistances genes - unobserved and subject to potential mutations that might affect their functions.

High throughput sequencing can address this by sequencing and assembling entire plasmids. Multiple institutions, including Applied Biological Materials Inc, seqWell, Massachusetts General Hospital Center for Computational and Integrative Biology, and CD Genomics, offer plasmid sequencing on Illumina platforms. While short read sequencing is accurate, the turnaround time is less than ideal for quick validations (>2 weeks) and the reliance on fragmenting before sequencing could prevent complete assembly - especially if the plasmid contains repetitive sequences much longer than Illumina read length. Indeed, some of the previously mentioned institutions charge a fee for assembling and annotating a plasmid sample and still cannot guarantee complete assemblies.

To address this shortcoming, several sequencing providers have turned to long read sequencing to ensure they capture full length plasmids; CD Genomics offers sequencing with Pacific Biosciences (PacBio) SMRT technology specifically for large plasmids, while Plasmidsaurus sequences plasmids exclusively with Oxford Nanopore Technologies (ONT). These facilities offer a quicker turnaround time and, in the case of Plasmidsaurus, a very low price (Table 1).

**Table 1.** Commercial Plasmid Sequencing Statistics. *additional cost for assembly and annotation **dependent on plasmid size. *** per well in 96 well plate. (Data collected Jan 2024)

| Company | Technology | Input Requirements | Price Per Sample | Turnaround Time |
| --- | --- | --- | --- | --- |
| ABM | Illumina | 1 - 50 ng | \$315* | 4-6 weeks |
| seqWell | Illumina | 600 - 1500 ng *** | NA | 2 weeks |
| MGH CCIB | Illumina | 1400 - 2275 ng | \$56.16 - \$70.20 | 5 - 8 business days |
| CD Genomics | Illumina/PacBio | 1400 - 2275 ng | \$270** | 15 business days |
| Plasmidsaurus | ONT | 300 - 2000 ng ** | \$15 - \$60** | 1 business day |

**Table 1:** R2C2 consensus read characteristics for different plasmids.

| Plasmid | Read type | Plasmid length (bp) | Median accuracy (%) | Average accuracy (%) |
| --- | --- | --- | --- | --- |
| tdTomato | R2C2 Consensus | 5,542 | 99.86 | 99.60 |
| pcDNA3.1 | R2C2 Consensus | 6,077 | 99.73 | 99.5 |
| pSpCas9 | R2C2 Consensus | 9,174 | 99.61 | 99.08 |
| lentiCRISPR | R2C2 Consensus | 14,874 | 99.54 | 98.48 |
| lentiCRISPR | ONT Subreads | 14,874 | 99.3 | 98.25 |

**Table 2:** Chopper generates highly accurate plasmid sequences. For each plasmid and run, the number of error-free sequences out of the 100 sequences produced by Chopper is shown.

| Sequencing Run | tdTomato | pcDNA3.1 | pSpCas9 | lentiCRISPR |
| --- | --- | --- | --- | --- |
| Individual | 99/100 | 100/100 | 85/100 | 100/100 |
| Pool | 100/100 | 100/100 | 47/100 | 100/100 |

Despite opportunities to outsource plasmid sequencing, there are several reasons why sequencing plasmids in house might be preferable to academic and commercial labs. These include using plasmid sequencing for training purposes, intellectual property concerns when outsourcing, sequencing plasmids that contain libraries of inserts, or simply the lack of availability of service providers.

ONT sequencing, using a MinION or P2 Solo, makes it possible for most labs to sequence complete plasmids in-house. There already exist protocols for the preparation and sequencing of plasmids [1] that perform very well but require the multiplexing of many plasmids to work and be economical.

Here, we set out to develop a method that could economically sequence individual or small pools of plasmids. We also wanted to develop an analysis tool to create error-free plasmid sequences.

To this end, we modified our R2C2 protocol [2] to use phi29-polymerase based rolling circle amplification (RCA) and T7 Endonuclease based debranching of the RCA product to generate linear dsDNA suitable for ONT sequencing. To reduce sequencing cost, we sequenced this dsDNA in our lab on an ONT P2 Solo using previously used flow cells and a modified library preparation protocol. We then processed the resulting data with a modified version of the established C3POa tool to parse individual sequencing reads and the newly developed Chopper to generate highly accurate polished full-length plasmid sequences.

## RESULTS

### Sequencing plasmids with a modified R2C2 workflow

We purchased four plasmids with known reference sequences from Addgene: pCSCMV:tdTomato (tdTomato), lentiCRISPR v2 (lentiCRISPR), pSpCas9(BB)-2A-Puro (PX459) V2.0 (pSpCas9), pcDNA3.1-GFP(1-10) (pcDNA3.1).

We then used a modified version of R2C2 to prepare each plasmid for sequencing - both individually and as part of a pool. Because the plasmids were already circular, we omitted the Gibson assembly step of R2C2 and instead directly amplified the plasmids with Rolling Circle Amplification (RCA) followed by debranching of the RCA product with T7 Endonuclease I. In this way, we produced long dsDNA containing multiple copies of the plasmids as tandem repeats (Fig. 1).

**Fig 1.**
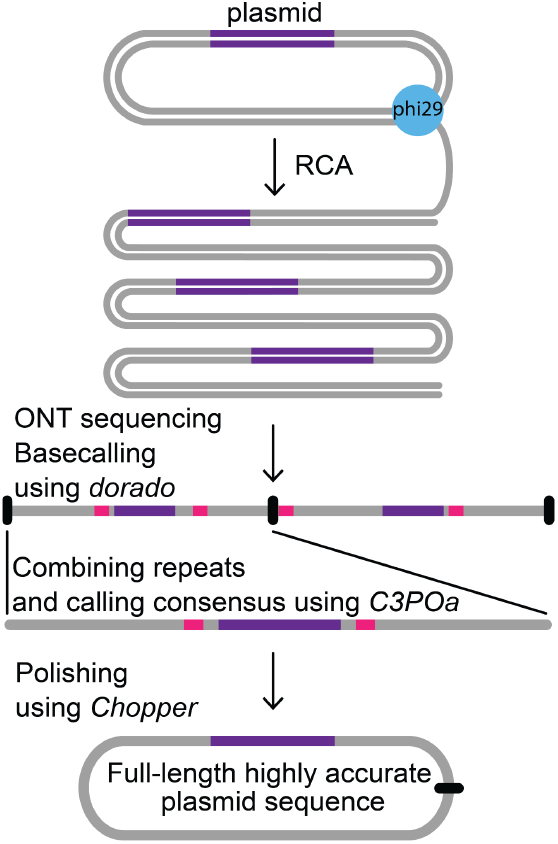
Plasmid processing, sequencing and analysis. Plasmids were processed with rolling circle amplification (RCA) to produce linear DNA containing multiple copies of the original plasmids. The linear DNA was prepped and sequenced on an ONT P2Solo. The resulting raw data was then processed into raw reads by dorado and consensus reads by an updated version of C3POa. Consensus reads were then combined into one highly accurate plasmid sequence for each plasmid using the newly developed Chopper.

After RCA and debranching, we performed a gel size selection of the resulting dsDNA, keeping only dsDNA that was over 10 kb in length. The size-selected dsDNA for each individual plasmid and the pool was then prepared for sequencing on an ONT P2 Solo using LSK114 ligation kits from ONT. Because this ONT library preparation represents the largest individual contributor to the cost of this plasmid sequencing workflow, we modified the protocol to use only 1/10th of the listed amount of sequencing adapter - the limiting reagent in the LSK114 kit.

For the actual sequencing runs we purposely sequenced on and in one case repeatedly reused PromethION flow cells that had been used for previous experiments and were at the end of their useful life. Because the accuracy of consensus sequences based on ONT reads plateaus after a few hundred raw reads [3], we reasoned that we would only need a few hundred reads per plasmid.

### Generating and Evaluating R2C2 Consensus reads

After sequencing each individual plasmid and the pool, we basecalled the reads with dorado (v0.7.3, model dna_r10.4.1_e8.2_400bps_sup@v5.0.0) and obtained 58,209 reads for tdTomato, 22,626 reads for pcDNA3.1, 136,012 reads for pSpCas9, 232,970 reads for lentiCRISPR, and 193,117 reads for the pool. The discrepancy in read numbers is related to the state of the flow cell before the experiment.

For further analysis, we initially wanted to use Tidehunter which recognizes tandem repeats in raw nanopore reads. However, Tidehunter [4] requires at least two full tandem repeats to be present in the raw read to detect it. This was a problem because the lentiCRISPR plasmid we aimed to sequence was about 15 kb long but only a small fraction of raw reads were longer than 20kb (Fig. 2) - shorter than the 30 kb it would take to cover two full repeats of this plasmid.

**Figure 2.**
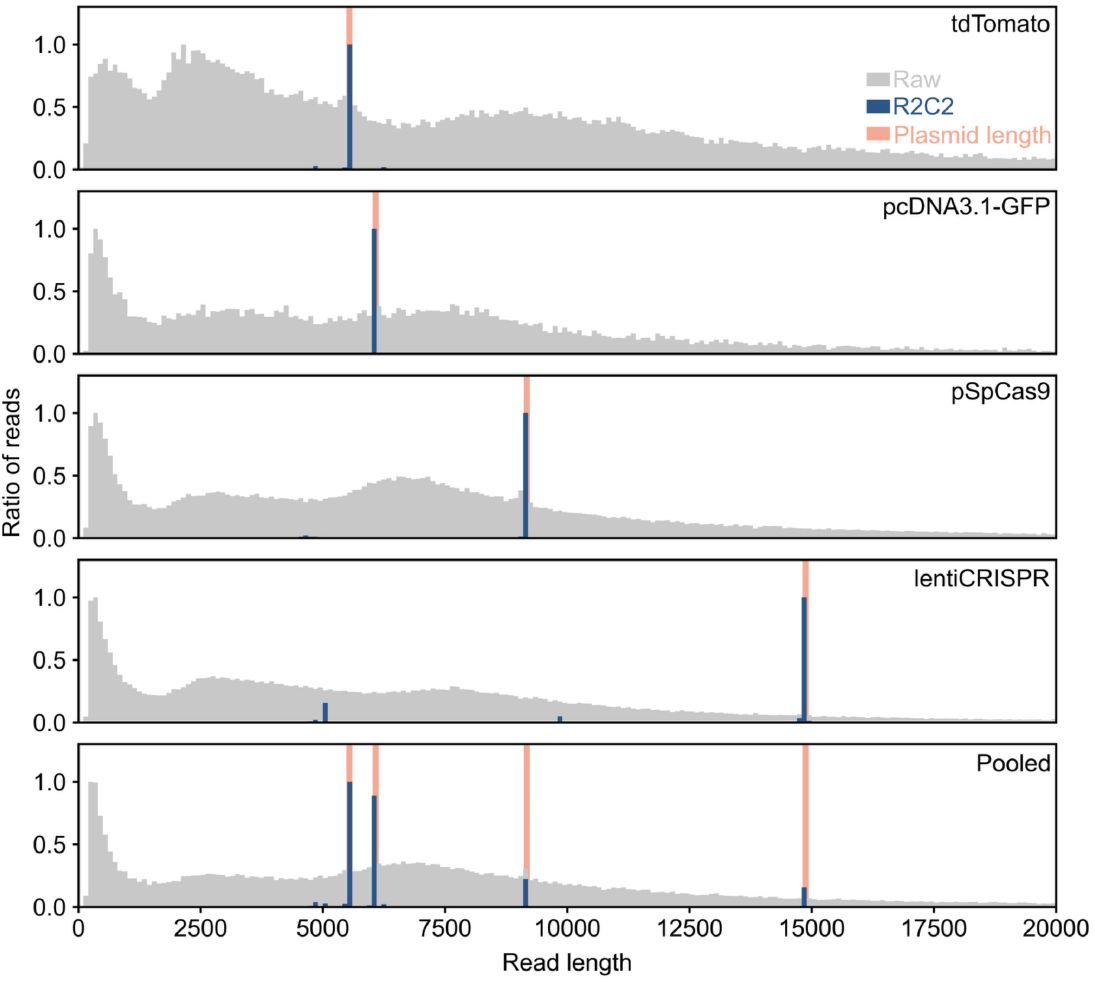
Sequencing plasmids with R2C2/C3POa. For each sequencing run, the read length distributions of the raw sequencing data (grey) and R2C2 consensus reads (blue) are shown as histograms. The length of sequenced plasmids (orange) included in each run is also shown.

We therefore chose to adapt C3POa, the standard R2C2 analysis pipeline to create consensus reads. As the C3POa tool was designed to create consensus reads from R2C2 molecules which have a specific structure it expects a known sequence of a DNA splint to mark repeats of input molecules. Since plasmids did not go through the full R2C2 workflow and therefore did not contain known splint sequences, we implemented a new option for C3POa: --nosplint or -ns, for raw data that does not contain a splint or other known sequence.

When --nosplint is set, instead of a user-defined known DNA splint sequence, C3POa uses a 200 bp anchor sequence taken from a few hundred bp into the raw read itself to identify repeat boundaries. During the development of this approach, we noticed that it can produce truncated plasmid sequences if there were internal repeats in a plasmid, as is the case for tdTomato. To prevent this C3POa skips raw reads where the distance between repeating anchor sequences was highly variable, which indicated the anchor was part of an internal repeat.

We then used this updated C3POa version with the --nosplint option on the raw reads of the different runs. This produced 37,617 consensus reads for tdTomato, 10,361 consensus reads for pcDNA3.1, 38,430 consensus reads for pSpCas9, 23,541 consensus reads for lentiCRISPR, and 65,971 consensus reads for the pool. In all cases the length of these consensus reads matched the length of the plasmid reference sequences, including the 15kb long lentiCRISPR plasmid (Fig. 2)

This showed that R2C2/C3POa can be used to generate full-length reads of plasmids from 5 to 15kb.

Once we had generated R2C2 consensus sequences from all datasets using the updated C3POa, we evaluated their accuracy by aligning the output of both tools to the Addgene reference using minimap2. Accuracy was calculated by dividing the number of matches by the sum of matches, mismatches, and indels within the read alignment. The accuracy of the R2C2 consensus reads reached 99.86% ∼(Q28) for tdTomato but declined with plasmid length approaching the raw ONT accuracy for lentiCRISPR (Table 1).

We assumed that the decrease in R2C2 consensus read accuracy could be attributed to subread coverage decreasing as plasmid length increases. To validate this assumption, we visualized read accuracy for each plasmid as swarmplots (Fig. 3). As expected, shorter plasmids showed more highly accurate high-coverage reads than the longer plasmids.

**Figure 3.**
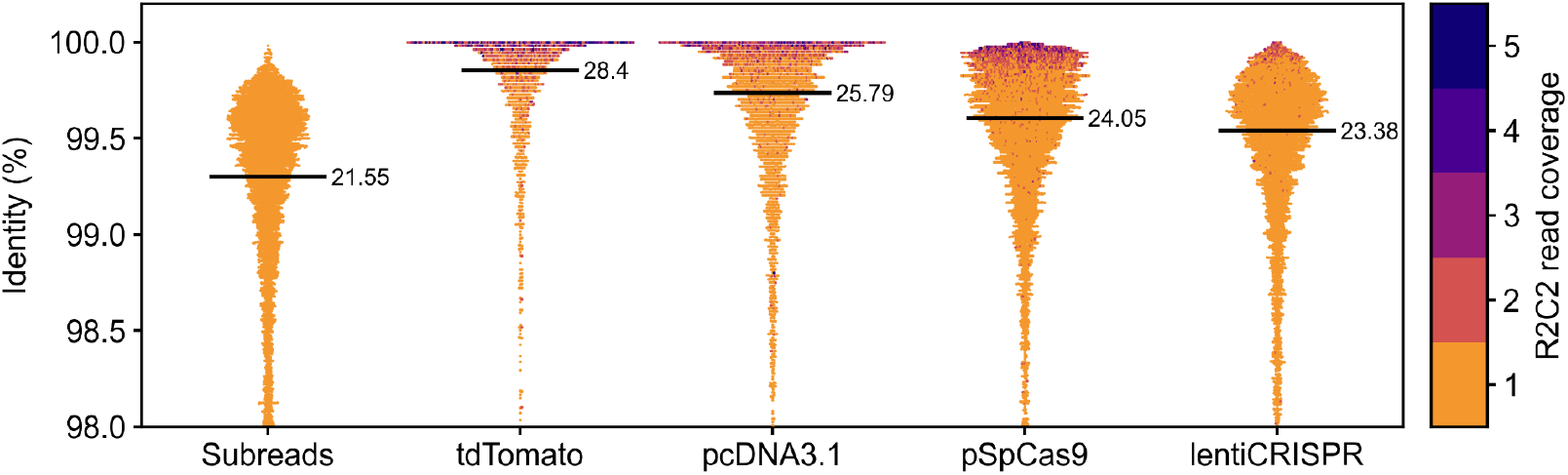
Evaluation of C3POa consensus accuracy. Accuracies of ONT subreads or R2C2 Consensus reads are shown for the indicated plasmids as swarmplots. Subreads were from the lentiCRISPR plasmid. Median read accuracy is shown as black bars with the exact Q score listed to its right.

In summary, using a modified C3POa tool, we processed R2C2 plasmid sequencing data to generate accurate full-length plasmid sequences.

### Generating highly accurate plasmid sequences with Chopper

After R2C2 consensus calling, we wanted to generate error-free sequences for each plasmid from both individual and pooled runs. We also wanted to take advantage of the medaka tool developed by ONT which is updated alongside the ONT *dorado* basecaller models and achieves very high accuracy.

To do so, we developed the Chopper tool (https://github.com/kschimke/Chopper/) (Fig. 4).

**Fig. 4:**
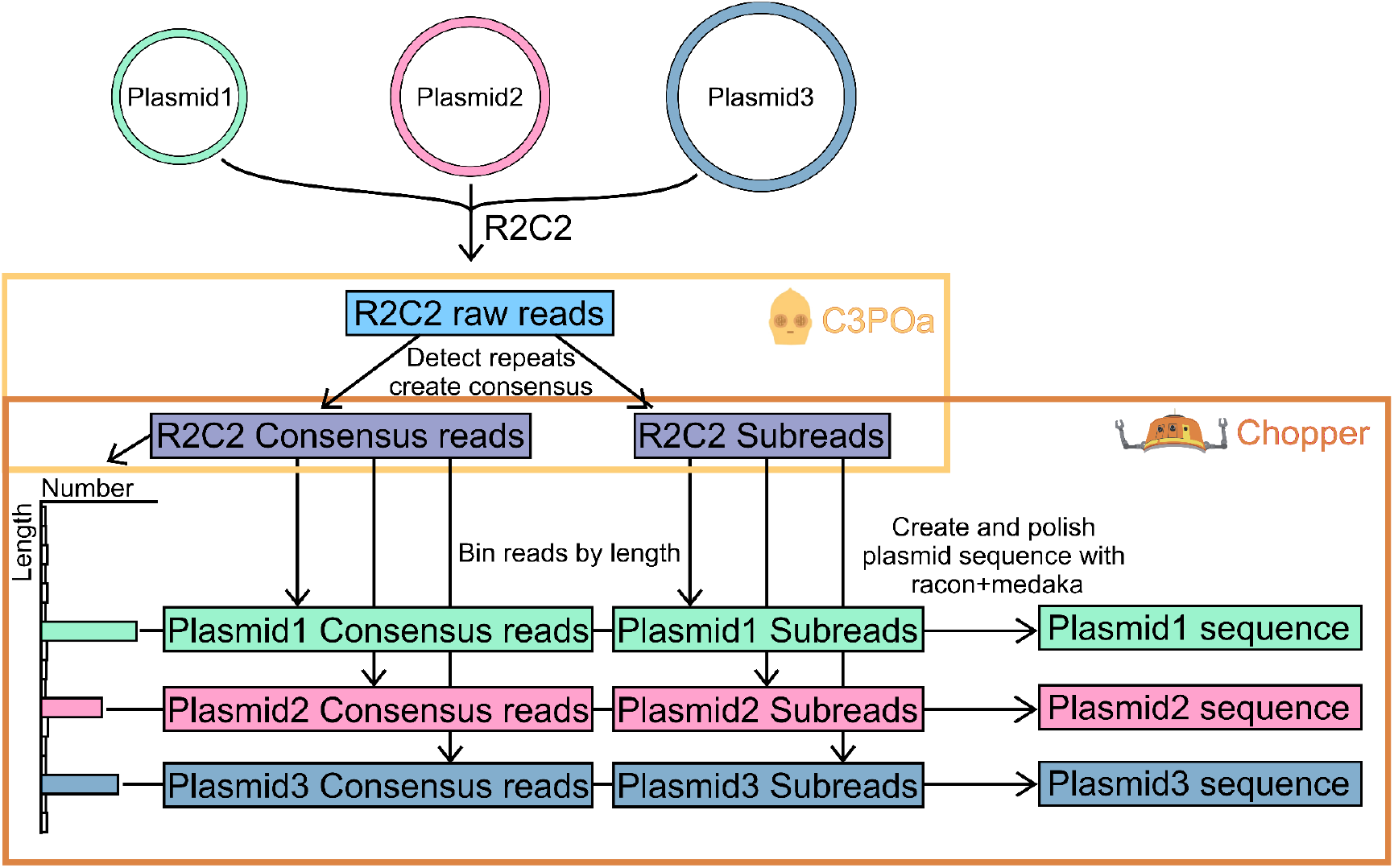
Chopper generates highly accurate plasmid sequences. Chopper accepts R2C2 consensus reads and their subreads produced by C3POa as input. It then bins R2C2 consensus reads by length to identify different plasmids in the sequencing data. For each length bin/plasmid, Chopper then generates a highly accurate plasmid sequence using racon and medaka for polishing.

Chopper bins R2C2 consensus reads by length and determines peak bins by comparing the size of the bins and finding local maxima. Chopper then separately processes the reads within each peak. It first creates a padded reference from the consensus read with highest coverage and raw Q score within the peak. Chopper then aligns the subreads with the highest Q scores to this reference using minimap2. Finally, Chopper uses racon [5] and medaka to polish the reference and then trims the polished sequence to represent a single full length plasmid.

We applied Chopper to the R2C2 consensus reads and subreads produced by C3POa for all five of our sequencing runs. To determine if the output of *Chopper* is dependent on the consensus read chosen to create the initial reference, we ran *Chopper* on the top 100 consensus reads for each length bin using the --iteration option in *Chopper*.

We evaluated the accuracy of the Chopper output sequences by aligning them to the plasmid references provided by Addgene with minimap2 [6] and used the same algorithm to calculate accuracy as we did for the R2C2 consensus reads. We then counted how many of the 100 Chopper iterations produced error-free plasmid sequences for each of the runs/plasmids

For the pcDNA3.1 and lentiCRISPR Chopper produced exclusively error-free sequences. For tdTomato 199/200 sequences Chopper produced from individual and pooled runs were error-free. However, Chopper struggled producing error-free sequences of pSpCas9 - 85/100 and 47/100 Chopper sequences for individual and pooled runs were error-free.

To investigate what caused Chopper to struggle with the pspCas9 plasmid, we determined where ONT subreads, R2C2 consensus, and Chopper sequences contained errors. By visualizing the alignments of reads from the pcDNA3.1 and lentiCRISPR runs to their respective plasmids, we found that errors were mostly random in ONT subreads, reduced in R2C2 consensus reads, and entirely absent in Chopper sequences.

For pSpCas9 however, ONT subreads, R2C2 consensus reads and Chopper read alignments all contained errors in two adjacent regions (Fig. 5A, center).

**Figure 5.**
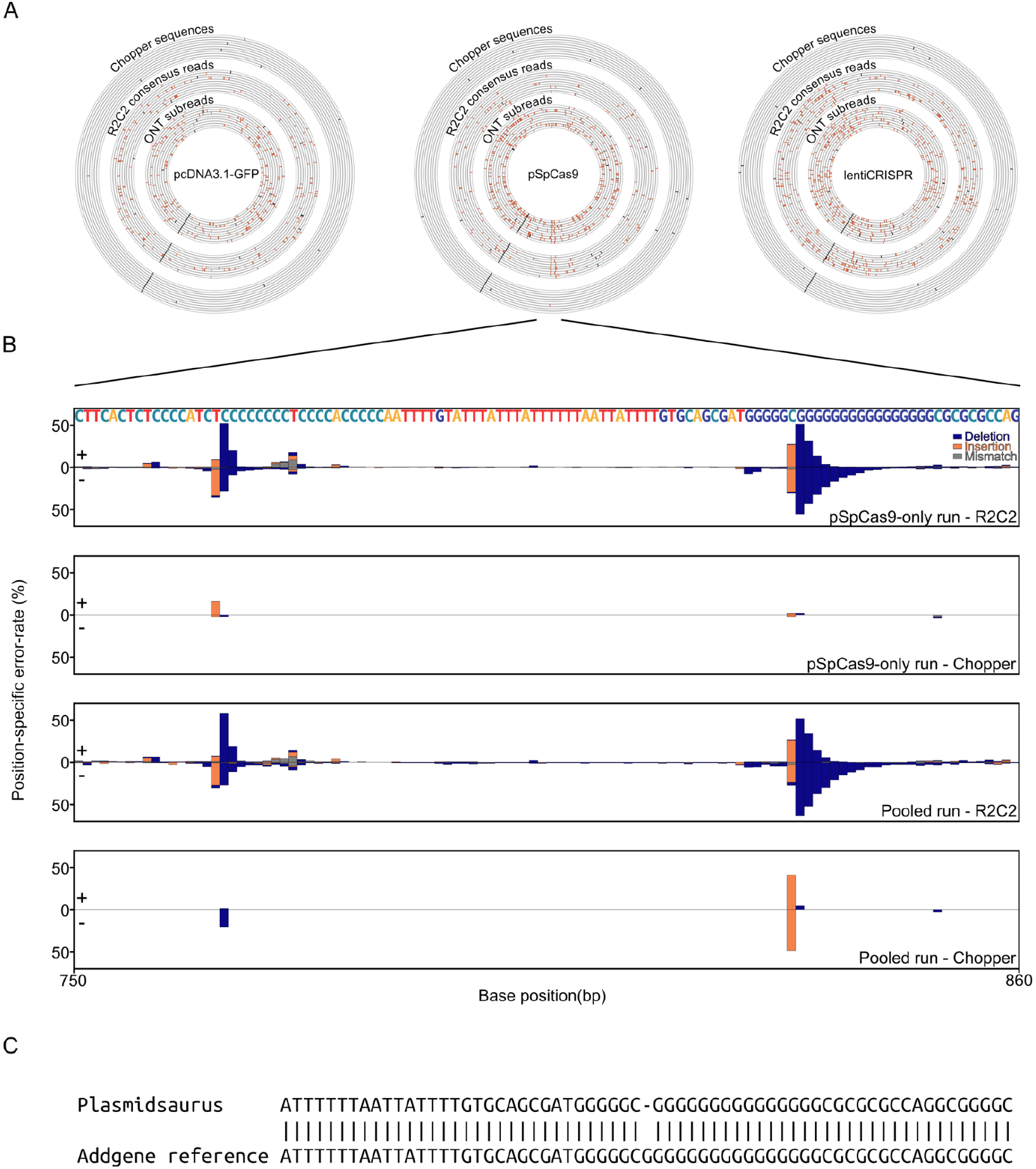
Chopper generates highly accurate plasmid sequences. ONT subreads, R2C2 consensus reads, and Chopper sequences are aligned to the indicated plasmid references. Mismatches and Indels and are indicated in red, alignment ends in black B). The error-rate of R2C2 consensus reads and Chopper sequences at specific positions of the pSpCas9 plasmid. These error-rates are shown as stacked bargraphs for a region of the plasmid that contains two homopolymers which represent a source for systematic sequencing error. C) Pairwise alignment of the pSpCas9 plasmid reference and the sequence Plasmidsaurus produced for that plasmid.

These regions contained two homopolymers of eight Cs and sixteen Gs, respectively (Fig. 5B). We quantified errors in R2C2 consensus reads across those two homopolymers and found a high level of systematic errors at those positions. Interestingly, the frequency of errors differed based on the direction of the reads. Further, while systematic errors appeared very similar between the individual and pooled sequencing run in the R2C2 consensus reads, Chopper sequences were much more likely to contain insertions at the G homopolymer in the pooled run.

This showed that small differences in sequencing runs can cause big differences in the calling of long homopolymers. To determine if the issues within the Chopper sequences were unique to Chopper or a problem with ONT sequencing technology in general, we sent all four plasmids in this study to Plasmidsaurus for sequencing and analysis. The resulting Plasmidsaurus assemblies of the tdTomato, pcDNA3.1, and lentiCRISPR plasmids were error-free, however the Plasmidsaurus assembly of the pSpCas9 plasmid contained an error in the longer G homopolymer (Fig. 5C).

This indicated that even a dedicated commercial sequencing provider with a likely bespoke assembly pipeline struggles to correctly assemble long homopolymer stretches, thereby highlighting the remaining limitations caused by highly systematic errors produced by ONT sequencing technologies.

## DISCUSSION

While it is difficult to argue with low cost and fast turnaround time for commercial whole plasmid sequencing, different labs might have different reasons to perform plasmid sequencing in-house.

Here, we have shown that a modified R2C2 protocol can be used to prepare individual or pools of plasmids for sequencing on ONT flow cells. We processed plasmids 5-15kb in length with this method and sequenced them on previously used flow cells for a limited time. In this way, we generated tens to hundreds of thousands of R2C2 reads covering full-length plasmids with accuracies up to 99.86%. Investigating these individual R2C2 reads might be of use to labs screening plasmid pools where each plasmid molecule might contain a (slightly) different sequence. An example of this might be plasmids containing a library of CRISPR gRNA sequences.

However, most investigators will likely be interested in a single error-free full-length plasmid sequence. To provide these single sequences, we developed Chopper, which can produce plasmid sequences that - with the exception of very long homopolymers - are error-free.

In addition to accuracy, time and cost will be considerations for investigators. The modified R2C2 protocol, sequencing, and data analysis took 2-3 days from sample to polished, complete plasmid sequence which was on par with the time required for outsourcing whole plasmid sequencing to Plasmidsaurus. We reduced the cost of the protocol by decreasing the volume of key ONT library preparation reagents used for each sample and negated the cost of a flow cell by sequencing on previously used flow cells. An additional factor in cost and time-savings is the ability of the R2C2/C3POa/Chopper workflow to handle pools of plasmids - as long as the plasmids have different lengths.

In summary, we present a combination of wet- and dry-lab tools that enable the fast, affordable, and highly accurate sequencing of plasmids.

## METHODS

### Library Preparation

We purchased four plasmids with known references from *Addgene*: pCSCMV:tdTomato, lentiCRISPR v2, pSpCas9(BB)-2A-Puro (PAX459) V2.0, pcDNA3.1-GFP(1-10). We pooled 125 ng of each plasmid to create a pool before R2C2 conversion. R2C2 was performed as in [3] with the exception of skipping the circularization step. In short, the individual plasmids and plasmid pool were amplified with rolling circle amplification (RCA) using Phi29 polymerase (NEB) and random hexamer primers (Thermo).

We performed 5 reactions for each plasmid and the pool using 100 ng of input each: [5 uL Phi29 Buffer (10X,NEB, M0269S), 1 uL Phi29 Polymerase(NEB, M0269S), 2.5 uL dNTP (10 mM;NEB N0447S), 2.5 uL Random hexamer primers (10 uM, Thermo, SO181), 10 uL plasmid (10ng/ul), 29 uL ultra-pure water]. RCA was incubated at 30°C for 16h. The RCA product was debranched by adding 2ul of T7 endonuclease I (NEB, M0302S) and incubating at 37°C for 2h. The resulting debranched DNA was then pooled and cleaned using a Monarch PCR and DNA Cleanup Kit (NEB). The purified RCA product was size-selected on an agarose gel: DNA at 10 kb and above was excised from the gel. R2C2 DNA was extracted from gel fragments using a Monarch DNA Gel Extraction Kit (NEB).

ONT libraries were prepared from R2C2 DNA using the ONT ligation sequencing kit V14 (ONT SQK-LSK114) following the manufacturer’s protocol while reducing the amount of Ligation Adapter to 1/10th of the recommended amount (0.5ul). The libraries were then sequenced on an ONT PromethION flow cell (R10.4.1) on a P2Solo for about 1 hour. Each flow cell was previously used for other experiments and reloaded with the plasmid R2C2 DNA after nuclease flush.

### Analysis

Raw nanopore sequencing data in the POD5 file format were basecalled using the *dorado* (v0.7.3) basecaller with the “sup” setting which uses the most accurate basecalling model (dna_r10.4.1_e8.2_400bps_sup@v5.0.0) to generate FASTQ files. FASTQ were then processed with C3POa (v3.2) to generate R2C2 consensus reads and R2C2 subreads.R2C2 consensus and subreads produced by individual and pooled plasmid runs were then processed by Chopper to generate 100 plasmid sequences for each size bin (−i 100). Cutoff percent for peaks (−t) was set to 10 for individual and and 2 pooled runs.

Alignments were performed using BLAT for the circular plots in Fig. 5 and minimap2 for everything else. BLAT [7] was chosen for the circular plots because it did a better job at aligning sequences to the reference in cases where a secondary alignment would be very short. These short secondary alignments were due to the circular nature of plasmids and the random start of sequencing reads within those plasmids.

## Data availability

R2C2 raw reads produced on the ONT P2 Solo are available under Bioproject PRJNA1209421 for the four individual plasmids and the pooled plasmids.

### Code availability

C3POa is available on GitHub at https://github.com/christopher-vollmers/C3POa. Chopper is available on GitHub at https://github.com/kschimke/Chopper.

